# Early rise and persistent inhibition of electromyography during failed stopping

**DOI:** 10.1101/2023.01.09.523332

**Authors:** Mitchell Fisher, Hoa Trinh, Jessica O’Neill, Ian Greenhouse

## Abstract

Reactively canceling movements is a vital feature of the motor system to ensure safety. This behavior can be studied in the laboratory using the stop signal task. There remains ambiguity about whether a “point-of-no-return” exists, after which a response cannot be aborted. A separate question concerns whether motor system inhibition associated with attempted stopping persists when stopping is unsuccessful. We address these two questions using electromyography (EMG) in two stop signal task experiments. Experiment 1 (n = 24) involved simple right and left index finger responses in separate task blocks. Experiment 2 (n = 28) involved a response choice between the right index and pinky fingers. To evaluate the approximate point-of-no-return, we measured EMG in responding fingers during the 100 ms preceding the stop signal and observed significantly greater EMG amplitudes during failed than successful stop trials in both experiments. Thus, EMG differentiated failed from successful stopping prior to the stop signal, regardless of whether there was a response choice. To address whether motor inhibition persists after failed stopping, we assessed EMG peak-to-offset durations and slopes (i.e., the rate of EMG decline) for go, failed stop, and successful stop (partial response EMG) trials. EMG peak-to-offset was shorter and steeper in failed stop trials compared to go and successful stop partial response EMG trials, suggesting motor inhibition persists even when failing to stop. These findings indicate EMG is sensitive to a point at which participants can no longer successfully stop an ongoing movement and suggest the peak-to-offset time of response-related EMG activity during failed stopping reflects stopping-related inhibition.

## INTRODUCTION

The ability to cancel an initiated movement is an essential aspect of daily life which allows individuals to adapt to dynamic environments. A common example is quickly stopping oneself from stepping into the street when a car unexpectedly appears. Stopping ongoing movements in response to an unexpected stimulus, often referred to as reactive inhibition, is one kind of response inhibition which can be studied in the laboratory with stop-signal tasks (Logan and Cowan, 1984; Verbruggen et al., 2019). During each trial of a stop-signal task, the participant rapidly responds to a go stimulus, however on a subset of trials, a stop signal is presented, and the participant attempts to abort their initiated movement. Behavioral data from the stop-signal task can be used to calculate a latent variable referred to as the Stop Signal Reaction Time (SSRT) which is an estimate of an individual’s average speed of stopping (Verbruggen et al., 2019). SSRT is a useful behavioral metric but is limited as it is an indirect measure and cannot be derived on an individual trial basis. The calculation of SSRT depends on the assumption of a horse race between competing independent Go and Stop processes, and whichever process finishes the race first determines the behavioral outcome. This has been a practical approach and has raised many important questions about the timing and mechanisms involved in response inhibition.

Stop-signal tasks are frequently paired with electromyography (EMG) which offers temporally precise measurements of muscle activity, a valuable supplement to behavioral data. Previous studies have examined EMG markers associated with stopping for a variety of reasons. De Jong et al. (1990), used this method to probe whether there was a ‘point of no return’ at which speeded, stimulus-driven movements could not be reactively aborted. This work demonstrated that partial EMG responses, i.e., EMG bursts in the absence of a button press, are a valuable measure of motoric inhibitory influence on trials in which participants successfully abort their response. These “partial responses” during successful stop trials reveal the point at which the Stop process overtakes the Go process at the level of the muscle and are only detectable with EMG since button presses are not registered on these types of trials. However, recent behavioral evidence (Du et al., 2022) contests the traditional horse race model, which posits that the Stop process is faster than the Go process. By suggesting Go and Stop processes take more or less equivalent amounts of time to complete, these newer findings call into question whether a ‘point of no return’ might be reached prior to the initiation of the stop process. Instead, the factor that determines successful versus failed stopping is the decision to delay or not delay the initiation of the Go response (Verbruggen & Logan, 2009). Delaying response initiation manifests as slowing when participants anticipate the possible presentation of a stop signal. These conflicting perspectives motivate a revisitation of the question whether a ‘point of no return’ exists through a careful examination of both behavior and EMG.

Others have investigated the timing and duration of the influence of inhibition within a responding effector at the single trial level through examination of partial response EMG during successful stop trials (Raud et al., 2020; Raud & Huster, 2017; Jana et al., 2020). A robust, single-trial electrophysiological marker of inhibition is valuable because traditional SSRT calculations, based solely on behavior, likely overestimate the speed of stopping by as much as 100 ms (Bissett et al., 2021; Skippen et al., 2019). The overestimation of the duration of the stopping process based on behavior alone may be due to the presence of ‘trigger failures’ when a participant fails to discern and respond to a stop signal meaning the stopping process was never ‘triggered’ (Matzke, Love & Heathcote, 2017; Matzke et al., 2019). In terms of the horse race model, the go process on such trials is executed uncontested despite the presentation of a stop signal. This also raises questions about the existence and ability to determine a possible ‘point of no return.’ However, EMG evidence supports the idea that stopping is faster than going. Raud & Huster (2017) observed participants initially displayed an EMG burst in response to the go cue, followed by a decrease in EMG 150 ms after the stop signal. It is possible this decrease in EMG is a result of the stop process finishing and canceling motor output (Raud et al., 2022). Notably, this occurred approximately 50 ms earlier than the SSRT estimation in that study and may be complemented by the engagement of antagonist muscles (Goonetilleke et al., 2010). Another recent study utilized a similar methodology to derive a single-trial estimate of stopping latency via EMG (Jana et al., 2020). This was accomplished by focusing on the time between the stop signal and the partial response EMG peak on successful stop trials as an estimate of the latency of stopping, building on previous examinations of partial response EMG activity (De Jong et al., 1990; McGarry et al., 2000).

Motivated by these recent investigations of EMG markers of stopping, we revisited the question of the ‘point of no return’ by more closely examining EMG onset timing at the individual trial level. We used simple and choice versions of the stop task to control for the possible influence of response selection. We predicted that a comparison of EMG activity prior to the stop signal would reveal a clear difference between failed stop and partial response (successful stop) trials. This finding would be useful in understanding the speed of stopping at the muscular level and bring context to previous work. In addition, we explored whether the time between peak EMG and EMG offset may reflect the operation of inhibitory processes acting at the level of the muscle. Specifically, we predicted a faster ‘ramp down’ of EMG activity on failed Stop trials than on Go trials. This finding would suggest Go and Stop processes both run to completion on failed stop trials, and the effect of inhibitory mechanisms may still be observable in muscle activity.

## METHODS

### Participants

Twenty-four participants (mean age = 23.7 years, std = 2.9 years, 10 female) completed simple response versions of a Go Task (GT) and a Stop Task (ST) in Experiment 1. Twenty-eight participants (mean age = 23.5 years, std = 4.1 years, 10 female) completed choice response versions of a GT and a ST in Experiment 2. Sample size was selected based on previous studies using similar methods (Raud et al., 2020; Raud & Huster, 2017; Jana et al., 2020). All participants were right-handed (self-reported) and provided written informed consent according to the institutional review board at the University of Oregon. All data and code are available from the Open Science Framework at this link: https://osf.io/9dp7x/

### Behavioral Tasks

Participants performed visual Go and Stop Tasks which were run in MATLAB using custom code (MathWorks). The tasks followed the recommendations set in Verbruggen et al. (2019), with the lone deviation that Experiment 1 did not contain a choice element. Participants were seated approximately 60cm from the screen and responded by pressing buttons that were interfaced with a MakeyMakey (JoyLabz, LLC) circuit board. Both experiments consisted of a GT and a ST (Figure 1A).

**Figure 1.**
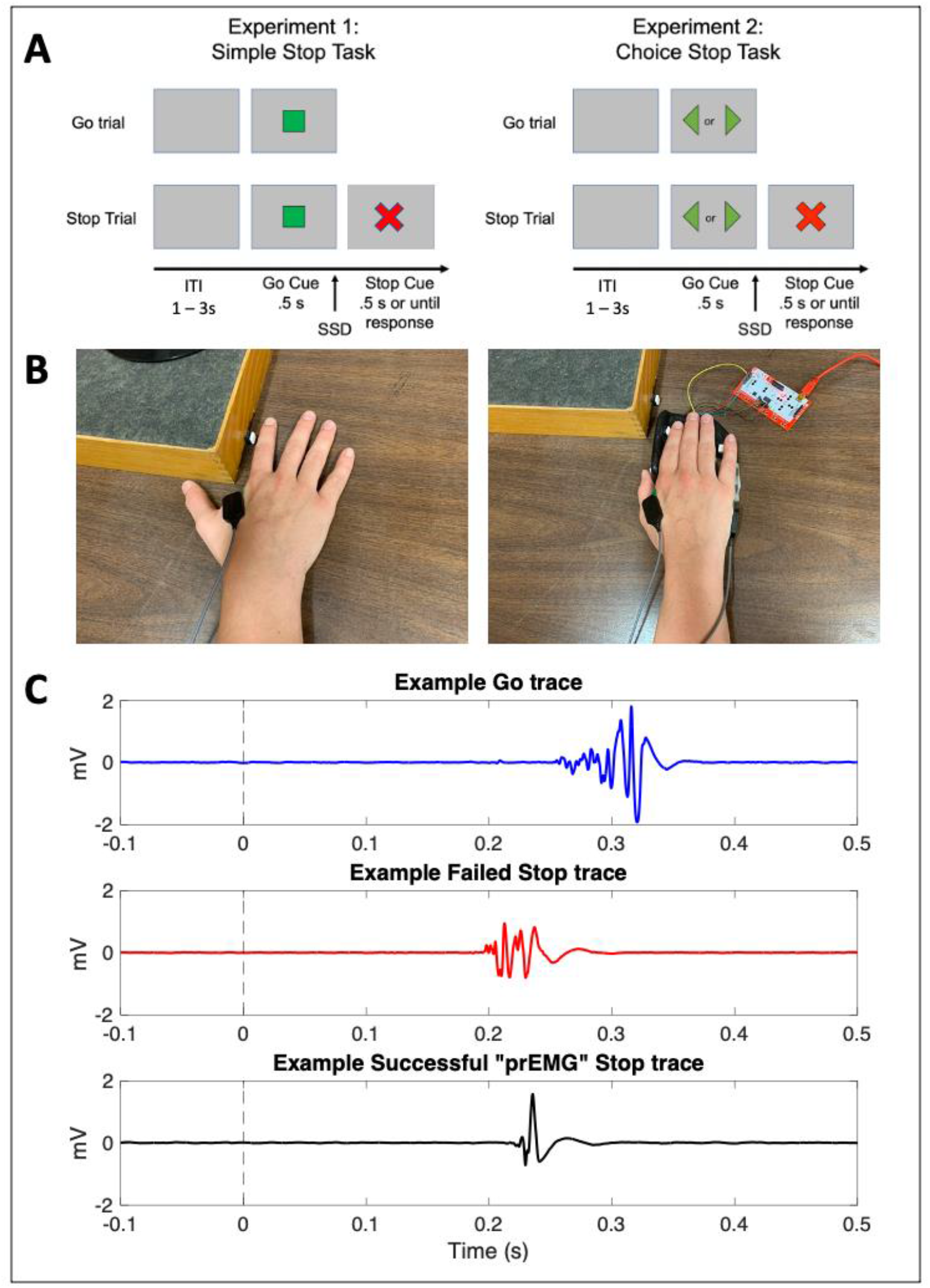
A) Visual task presentation in Experiments 1 and 2. B) Electrode placement and button configuration (left panel = Exp. 1, right panel = Exp. 2). C) Example EMG traces for 3 trial conditions that contain EMG responses: Go, Failed stop and Successful stop (prEMG), with time 0 indicating Go stimulus onset.

#### Experiment 1: Simple Go and Stop Task

Experiment 1 consisted of simple response tasks (no choice element) that participants performed with one hand and then the other in a randomized order. Participants’ hands were positioned palms down, shoulder-width apart on a table in front of the visual display. Responses were made by laterally moving the forefingers toward the midline of the body to depress a horizontally positioned button. Participants performed the GT with one hand, the GT with the other hand, the ST with the first hand, and then the ST with the second hand.

During the GT, participants completed at least 18 trials as practice prior to testing followed by one block of 30 test trials per hand. Each trial consisted of a blank screen inter-trial interval (ITI) of 1 to 3 s (random, uniform distribution) followed by the presentation of a Go signal. The Go signal on each trial was a green square presented at the center of the screen, and participants were instructed to respond as quickly as possible by depressing the response button. The Go signal remained on the screen for 0.5 s and a correct response could be registered for 1 s from Go signal onset. No feedback was provided to participants during the GT. Average GT reaction time (RT) was used to determine the RT threshold for performance feedback during the ST as described below.

In the ST, participants were informed that each trial would begin with a Go signal as described above and that on some trials, this would be followed at a short interval by a Stop signal (a red ‘X’) that replaced the Go signal. Trial duration, ITI, and jitter were the same as the GT. The Go stimulus was presented for 0.5 s, and on stop trials, the Stop signal lingered for 0.5 s. Participants were also familiarized with the ST by performing at least 18 practice trials, until they felt comfortable with the task. During testing, participants completed 108 trials total with each hand, consisting of 72 go trials (66%) and 36 stop trials (33%). The stop signal was presented at a variable Stop Signal Delay (SSD) using three independent staircases (starting at 40, 80 and 120 ms) that were adjusted according to participant performance. The SSD increased by 20 ms if stopping was successful and decreased by 20 ms if stopping was unsuccessful. The task was broken up into 4 blocks with 27 trials in a block. Between blocks, participants were required to take at least a 90s break to prevent fatigue.

Participants were instructed to not delay their responses in anticipation of the stop signal. To enforce speeded responses, participants received a ‘Speed up’ cue on go trials when button press RT was greater than 2.5 standard deviations above the GT mean RT. Trials that resulted in the presentation of the ‘Speed up’ cue were not analyzed, and participants were excused from the study if they received greater than 20 ‘Speed up’ cues due to inadequate performance.

#### Experiment 2: Choice Go and Stop Task

The Go and Stop Tasks in Experiment 2 consisted of a choice between left and right responses, mapped to the index and pinky fingers of the right hand, respectively. Participants’ right hands were positioned on a custom-built response device with buttons positioned against the lateral portion of the distal phalanx of the index finger and beneath the tip of the pinky finger. Index finger responses were the same as the right index responses in Experiment 1 involving a lateral movement toward the midline of the body. Pinky finger responses involved a downward button press (see Figure 1B right panel).

The GT consisted of two blocks of 30 trials (60 trials total) with an equal number of left and right responses randomly distributed across trials. Participants completed at least 18 trials of practice prior to testing. The ITI was 1.2 s and each trial was jittered from 1 to 3 s (random, uniform distribution). The Go signal was either a right-or left-pointing green arrow for the index and pinky responses, respectively, and was displayed for 0.5 s. Correct responses could be registered up to 1 s from Go signal onset. As in Experiment 1, no feedback was presented during the GT, but the mean GT RT was used to determine the threshold for presenting feedback during ST performance.

The ST was similar to the GT except it consisted of eight blocks of 27 trials (216 trials total). One third of the trials (72 trials) were stop trials, with 36 stop trials per finger. The stop trials were distributed randomly across the experiment. On each stop trial, the Go stimulus was replaced with a stop signal (red ‘X’) at a variable SSD. As in Experiment 1, the SSDs were adjusted dynamically based on performance with 3 staircases for each response finger (six independent staircases total). The staircases started at 100, 125, and 150 ms and increased or decreased by 20 ms following successful or failed stop trials, respectively. Participants received ‘Speed up’ feedback on go trials during the ST when they failed to respond within 2 standard deviations of their mean Go RT determined from the GT. As above, if participants received 20 ‘Speed up’ cues, the experiment would end, and the data was excluded from analysis. Participants also received ‘Try to stop’ feedback on stop trials when they failed to cancel their movement.

#### EMG Procedure

EMG was recorded using bipolar surface electrodes connected to an eight-channel Bagnoli (Delsys) amplifier (amplification x1000, bandpass filtered 50-450 Hz). EMG was sampled at 5000 Hz and recorded using the VETA toolbox (Jackson & Greenhouse, 2019).

In Experiment 1, EMG electrodes were placed over the left and right first dorsal interossei (FDI) muscles. In Experiment 2, EMG electrodes were placed over the left and right FDI and the right abductor digiti minimi (ADM) muscle. A 4 s EMG sweep was recorded for each trial of the task and visualized by the experimenter on a separate screen adjacent to the stimulus presentation monitor. During the practice trials, participants were coached to keep their responding fingers close to the response buttons and to keep the EMG signal uncontaminated by noise when not responding to improve sensitivity to EMG events.

In both experiments, participants were instructed to maintain a tonic contraction of the FDI muscle of the non-responding hand throughout task performance. Prior to testing, the maximum voluntary contraction (MVC) of the FDI in the non-responding hand was measured, and participants were trained to maintain a tonic contraction of approximately 10% MVC. Online feedback displayed individualized reference lines on the EMG recording figure corresponding to 10% of the MVC, and helped ensure that participants maintained a stable contraction. We consider the possible influence of these contractions on the behavioral performance and EMG in the responding hand but do not discuss the tonic EMG further in the current manuscript.

### Statistical Analyses

#### Behavioral Measures

Behavioral metrics of interest were calculated for each task condition in both experiments. These included means and standard deviations of button Go RT, mean SSD, mean stopping accuracy (%), integration SSRT (Verbruggen et al., 2020), and means and standard deviations of Failed stop button RT.

#### EMG Analysis

EMG data were analyzed offline using the VETA toolbox (Jackson & Greenhouse, 2019) in Matlab 2019a (Mathworks) in two stages. First, EMG bursts were identified using an automatic detection algorithm. Second, data from each trial were visualized, and manual corrections were made where necessary. This included rejection of sweeps contaminated with artifacts and adjustments to mark the onsets and offsets of EMG bursts. EMG data were smoothed using a root mean squared method (‘movRMS’ via the DSP system Toolbox) with a sliding window of 12 sample points, equivalent to 2.4 ms and rectified.

EMG metrics of interest included means and standard deviations of EMG burst onset, peak, and offset times. These were calculated separately for Successful go trials (Go EMG RT, Go EMG peak, Go EMG offset), Failed stop trials (Failed EMG RT, Failed EMG peak, Failed EMG offset), and Partial responses on successful stop trials (Partial EMG RT, Partial EMG peak, Partial EMG offset). Additionally, EMG onset-to-peak times, and EMG peak-to-offset times for all EMG bursts were calculated for each trial type and task condition. Cancel time was calculated as time elapsed from stop signal onset to EMG peak for Partial responses (Jana et al., 2020).

In addition, the electromechanical delay (EMD), i.e., the time between EMG onset and button press, was calculated for Go and Failed Stop trials. EMG peak-to-offset times were calculated along with the EMG peak-to-offset slope for all response conditions. Slope was calculated using a linear fit of the mean EMG peak to the mean offset of the EMG burst for each response condition for each participant. Differences in the EMG peak-to-offset times across conditions could be driven by especially slow Go responses. Therefore, we performed a separate analysis using EMG peak-to-offset times for Go trials with button RTs shorter than the slowest Failed stop RT for each participant, matching the maximum allowable button RT across these two conditions.

#### Statistical Tests

Statistical analyses were performed in Jamovi (version 2.2, Jamovi, 2021) except for Bayesian tests which were performed in JASP (version 0.14.1, JASP, 2020). For Experiment 1, a 2 × 3 repeated-measures ANOVA with the factors Hand (Left, Right) and Trial Type (GT Go, ST Go, Failed Stop) tested for differences in button RTs and EMD. Bayesian paired *t*-tests tested for equivalence of performance for SSD, stop accuracy and SSRT between hands (Experiment 1) and response choices (Experiment 2), and Bayes Factors (BF) were derived.

A 2 × 4 repeated-measures ANOVA with the factors Hand (Left, Right) and Trial Type (GT Go, ST Go, Failed Stop, Partial Response) tested for differences in our EMG metrics of interest. A significant main effect of trial type would indicate that EMG activity differs across the response conditions. Here, we hypothesized that partial responses would show a later onset and possibly more gradual ramp up than the other response conditions, based on the assumption that the likelihood of stopping increases with slower responses. Moreover, we hypothesized that there would be differences in the ramp down in EMG activity between Go and Stop trials, both successful partial responses and failed responses.

The same statistical tests were conducted for Experiment 2 with the factor Hand replaced by the factor Finger (Index, Pinky). Post-hoc follow up tests were Bonferroni corrected.

To evaluate the point of no return, i.e., the point at which a button press cannot be aborted, mean EMG amplitude for the 100 ms preceding stop signal onset was compared between failed and successful stopping. The rationale for this 100 ms window is based on the assumption that the go stimulus has been fully processed by this time, and EMG activity within this window is most likely associated with response initiation. Significantly greater mean EMG activity during failed than successful stopping in this time window would be consistent with the interpretation that the point of no return precedes stop signal onset.

The time between EMG peak and EMG offset provides a measure of the speed with which EMG activity resolves following the engagement of a muscle. If EMG activity decreases significantly faster on failed stop than go trials and resembles successful stop trials, this would suggest inhibition associated with stopping persists even when the stop process fails. In other words, the motor system is inhibited, but too late to prevent the completion of the go response. In this case, the rate of decline in EMG activity on failed stop trials would provide a novel marker of motor system inhibition.

## RESULTS

### Behavioral Data

#### Experiment 1

Behavioral data are presented in Table 1 for both experiments. Data for the right and left simple RT tasks were very closely matched. Analysis of button press RTs (GT Go, ST Go and ST failed stop trials) indicates no differences between hands [*F* (1,23) = .33, *p* = .57, *η*^2^ = .001] and a main effect of trial type [*F* (2,23) = 90.28, *p* < .001, *η*^2^ = .38]. Successful stop trials were not included in this analysis due to the absence of a button press. A Bonferroni-corrected 2 × 2 repeated measures ANOVA revealed no differences between hands [*F* (1,23) = .04, *p* = .85, *η*^2^ = .00] and significantly longer ST go trials compared with GT go trials [*F* (1,23) = 103.69, *p* < .001, *η*^2^ = .41]. This confirms participants slowed their responses on go trials in the context of the ST. Paired-samples *t*-tests indicated SSRT, SSD, and stopping accuracies did not differ between hands (all *p*’s > .05). Bayesian tests for equivalence between hands for SSRT, SSD, and stopping accuracies showed anecdotal evidence for all measures [all BF_01_ between 1.48 and 2.66].

**Table 1.** Means (standard deviations) for behavioral and EMG measures in Experiment 1 (n = 24) and 2 (n = 28). GT = go task, ST = stop task, RT = reaction time, prEMG = successful stop trials that included a partial EMG response, SSD = stop signal delay, SSRT = stop signal reaction time.

| Behavioral Measures | Experiment 1 |  | Experiment 2 |  |
| --- | --- | --- | --- | --- |
|  | (Simple Task) |  | (Choice Task) |  |
|  | Right<br>(FDI) | Left<br>(FDI) | Index<br>(FDI) | Pinky<br>(ADM) |
| GT RT (ms) | 352 (32) | 352 (30) | 454 (58) | 456 (59) |
| ST Go RT (ms) | 410 (41) | 408 (35) | 453 (42) | 459 (48) |
| ST Fail RT (ms) | 357 (33) | 353 (32) | 390 (31) | 396 (43) |
| GT EMG RT (ms) | 203 (23) | 207 (21) | 258 (30) | 271 (28) |
| ST Go EMG RT (ms) | 245 (34) | 246 (33) | 260 (30) | 281 (32) |
| ST Fail EMG RT (ms) | 208 (31) | 206 (26) | 223 (25) | 240 (26) |
| ST prEMG RT (ms) | 226 (33) | 222 (32) | 257 (29) | 278 (46) |
| SSRT (ms) | 245 (20) | 256 (29) | 233 (39) | 246 (41) |
| Cancel Time (ms) | 168 (27) | 166 (29) | 158 (32) | 161 (33) |
| SSD (ms) | 118 (23) | 113 (23) | 162 (27) | 167 (28) |
| Go trial accuracy (%) | 99.9 (0.3) | 99.8 (0.5) | 99.4 (1.4) | 99.2 (1.7) |
| Stop trial accuracy (%) | 66.1 (8.2) | 63.9 (9.5) | 65.1 (8.9) | 68.3 (10.8) |

#### Experiment 2

Generally, RTs between fingers (FDI and ADM) in Experiment 2, exhibited similar RT patterns as between hands in Experiment 1. However, unlike Experiment 1, no slowing was observed between button GT Go RT and the ST Go RT for either finger [*F* (1,27) = 1.20, *p* = .29, *η*^2^ = .00]. Failed stop trial button RTs were shorter than go trials in both fingers [*F* (1,27) = 64.3, *p* < .001, *η*^2^ = .281]. Paired-samples *t*-tests showed that SSRT and SSD were similar between fingers (all *p*s > .05), while stopping accuracy was higher for pinky responses than index finger responses (*p* < .05, *d =* .33). Bayesian tests for equivalence between hands for SSRT, SSD, accuracies showed anecdotal evidence for all measures [all BF_01_ between .35 and 2.89].

### EMG Data

#### Experiment 1: EMG Reaction Times and Electromechanical Delays (EMD)

EMG RTs were similar between hands [*F* (1,23) = .01, *p* = .94, *η*^2^ = .001] but varied across GT, ST Go, failed stop, and successful stop (partial EMG) response types [*F* (3,23) = 41.63, *p* < .001, *η*^2^ = .24]. Like the button-press results, slowing was observed for both hands for the ST go trials relative to the GT, and failed stop trials had the shortest latency. A Bonferroni-corrected 2 × 2 repeated measures ANOVA showed ST go trials were significantly slower than GT go trials [*F* (1,23) = 75.11, *p* < .001, *η*^2^ = .35].

Successful stopping was defined by stop trials that did not have a button response registered, however inspection of EMG data revealed that on approximately half of the stop trials (16.7 ± 4.7 trials per hand) EMG activity increased, indicating participants started to respond, but did not fully execute the button press. These trials are meaningful because there is an implied inhibitory process that prevents the completion of the button press response. The mean latency of EMG onsets for these partial response trials was directly in between the ST Go trial mean latency and the failed Stop trial mean latency for each hand (Table 1), consistent with the idea that even early EMG activity can be successfully suppressed by the stop process. No differences in EMDs were found between hands *[F* (1,23) = .49, *p* = .49, *η*^2^ = .003]. A main effect across trial type was observed for EMD [*F* (2,23) = 14.61, *p* < .001, *η*^2^ = .15] with ST go trials being longer than the other two trial types.

#### Experiment 1: Response EMG Profiles

Further inspection of the EMG profile of the responding finger was warranted to determine how partial response EMG RTs differed from failed stop EMG RTs and how they each compared with go trial responses (Figure 2). To do this, we identified all trials with EMG responses and measured time epochs including: onset of EMG activity to the peak of EMG activity, peak of EMG activity to EMG offset (return to baseline), and cancel time. Cancel time is a single trial estimate of stopping latency and is the time from the stop stimulus onset to the peak of EMG activity, as described by Jana et al. (2020). Cancel time is only appropriate to measure in successful stop trials, and as such was evaluated using paired-samples *t-*tests on prEMG trials. No difference was observed between hands (*p* = .79, *d =* .06).

**Figure 2.**
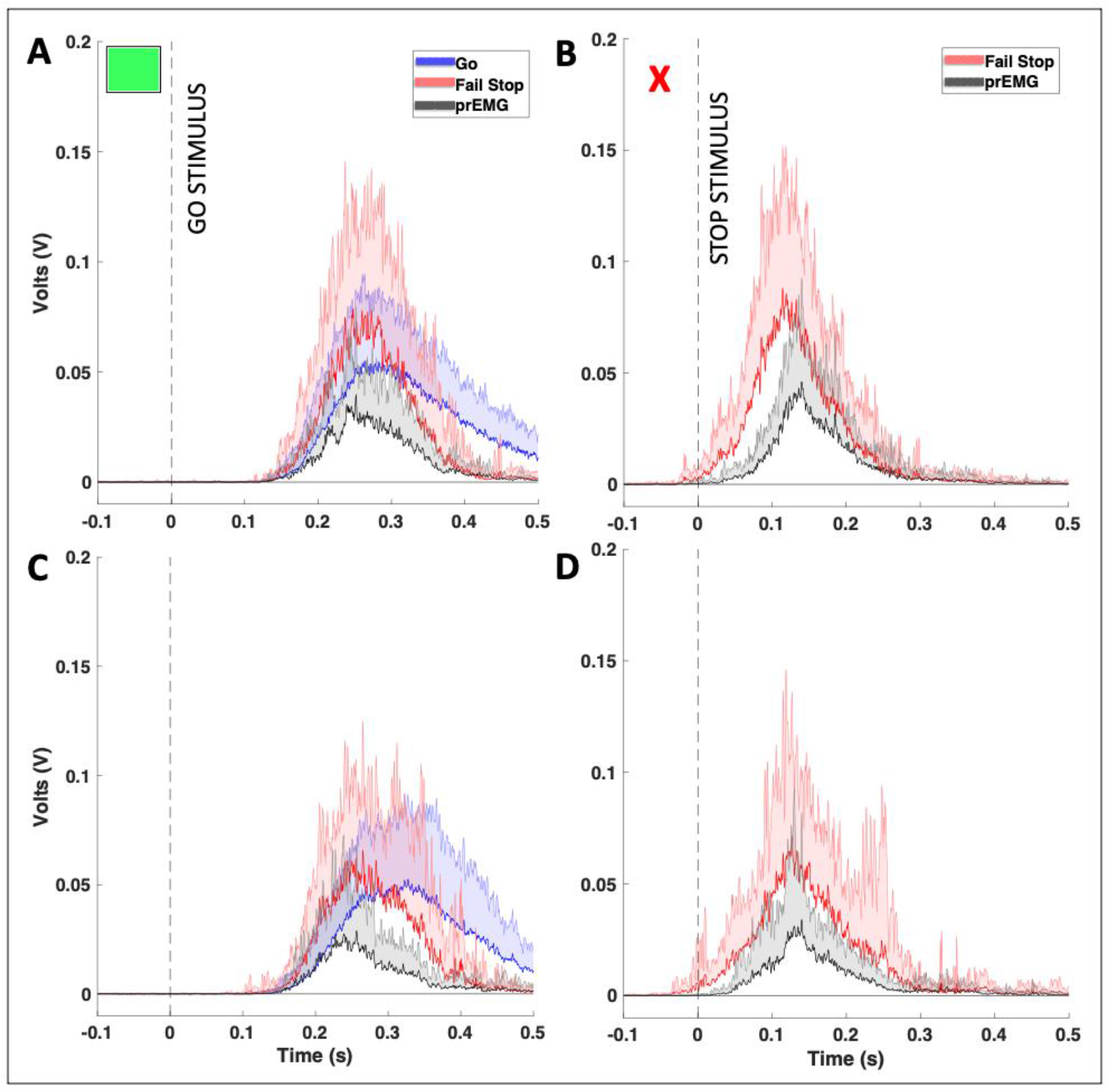
Experiment 1 (n=24) mean ± std response EMG aligned with Go stimulus (A & C) and Stop stimulus (B & D). Responses from the right hand are in A & B, while responses from the left hand are in C & D. Blue traces are from go trials, while black and red traces are from successful stop trials that had a partial response (prEMG) and failed stop trials, respectively. Data are smoothed via root mean square method and rectified.

Onset to peak times were similar between hands [*F* (1,23) = 3.04, *p* = .09, *η*^2^ = .008], but differed across conditions as expected [*F* (3,23) = 53.35, *p* < .001, *η*^2^ = .34], (Table 2). When observing peak EMG to EMG offset times (Figure 4 A & C), right hand values were significantly longer than left hand values [*F* (1,23) = 5.39, *p =* .029, *η*^2^ = .038]. Notably, differences in peak EMG to EMG offset times across conditions were also significant [*F* (3,23) = 14.4, *p* < .001, *η*^2^ = .148]. To evaluate whether the decline in EMG activity differed between failed stop and ST go trials, we performed a post-hoc 2 × 2 repeated measures ANOVA with Bonferroni correction. This test revealed a significant difference between ST go trials and failed stop trials [*F* (1,23) = 6.73, *p =* .016, *η*^2^ = .021] and no significant difference between hands [*F* (1,23) = 2.84, *p =* .106, *η*^2^ = .042]. Slope was also calculated to index the rate of decline from peak EMG to EMG offset for each hand and all conditions (Table 2). Failed stop trials exhibited a significantly steeper slope than ST go trials and prEMG trials [*F*(1,23) = 32.768, *p* < .001, *η*^2^ = .259], and there was a significant interaction between hands and trial type [*F*(3,69) = 3.658, *p =* .034, *η*^2^ = .014]. To evaluate the significant interaction effect, a 2 (Go vs Fail) × 2 (Left vs Right) repeated measures ANOVA showed that the slope difference between Go and Failed stop trials in the right hand was greater than in the left hand [*F*(1,23) = 4.79, *p =* .039, *η*^2^ = .017]. These results indicate EMG burst activity declined more quickly on failed stop trials than go trials, despite the failure to stop, a pattern consistent with lingering inhibition on failed stop trials.

**Table 2.** Means (standard deviations) for derived EMG measures in Experiment 1 (n = 24) and Experiment 2 (n = 28). EMD = electromechanical delay, prEMG = successful stop trials that included a partial EMG response.

| Derived EMG Measures | Experiment 1 |  | Experiment 2 |  |
| --- | --- | --- | --- | --- |
|  | (Simple Task) |  | (Choice Task) |  |
|  |  | Left | Index | Pinky |
|  | Right (FDI) | (FDI) | (FDI) | (ADM) |
| GT Go EMD (ms) | 149 (22) | 145 (23) | 195 (46) | 186 (43) |
| ST Go EMD (ms) | 164 (27) | 162 (23) | 193 (29) | 177 (36) |
| ST Fail EMD (ms) | 150 (23) | 148 (19) | 165 (27) | 156 (29) |
| GT Onset to Peak (ms) | 82 (24) | 83 (25) | 117 (48) | 101 (34) |
| ST Go Onset to Peak (ms) | 88 (24) | 95 (23) | 108 (24) | 95 (28) |
| ST Fail Onset to Peak (ms) | 65 (18) | 72 (17) | 95 (18) | 86 (24) |
| ST prEMG Onset to Peak (ms) | 52 (14) | 72 (17) | 92 (18) | 85 (22) |
| GT Peak to Offset (ms) | 107 (54) | 84 (41) | 109 (52) | 105 (37) |
| ST Go Peak to Offset (ms) | 92 (45) | 74 (21) | 116 (56) | 92 (33) |
| ST Fail Peak to Offset (ms) | 78 (25) | 65 (27) | 104 (49) | 84 (29) |
| ST prEMG Peak to Offset (ms) | 58 (27) | 48 (18) | 100 (46) | 80 (26) |
| ST Go EMG Offset Slope (mV/s) | -0.6 (0.3) | -0.7 (0.5) | -0.7 (0.4) | -0.4 (0.3) |
| ST Fail EMG Offset Slope (mV/s) | -2.3 (1.7) | -1.7 (1.3) | -1.7 (1.1) | -1.2 (0.8) |
| ST prEMG Offset Slope (mV/s) | -1.1 (0.8) | -1.0 (0.7) | -0.8 (0.6) | -0.6 (0.5) |

Peak EMG values showed no differences between hands [*F* (1,23) = 2.94, *p =* .10, *η*^2^ = .025], but there was a significant difference across conditions [*F* (3,23) = 63.52, *p* < .001, *η*^2^ = .184]. In the right hand, the largest to smallest peak EMG values (mean ± sd) were GT go trials (1.80 ± 0.74V), ST go trials (1.80 ± 0.70V), failed stop trials (1.64 ± 0.70V) and finally, prEMG trials (0.97 ± 0.51V). The left hand exhibited a relatively similar pattern with the exception that go trials in the ST achieved the highest peak EMG values (1.55 ± 0.87V), followed by GT go trials (1.49 ± 0.78V), failed stop trials (1.39 ± 0.77V) and prEMG (0.69 ± 0.49V).

Finally, we investigated the EMG epoch 100 ms prior to the stop stimulus for failed Stop trials and successful Stop trials with prEMG to determine if the EMG activity in this time window differentiated between successful and failed stopping (Figure 2 B & D). We compared the mean rectified EMG signal within this time window between the two conditions with a paired-samples *t*-test. EMG activity in this period was significantly greater in failed stop trials compared to successful stop trials in the right (*p <* .05, *d* = .57) and left hands (*p =* .05, *d* = .42).

#### Experiment 2: EMG Reaction Times and Electromechanical Delay (EMD)

In experiment 2, EMG RTs were evaluated in a similar manner as above and raw values can be seen in Table 1. EMG RTs differed between fingers [*F* (1,27) = 28.805, *p* < .001, *η*^2^ = .063] as well as between conditions (GT go trials, ST go trials, failed stop trials, and successful stop trials with prEMG) [*F* (3,27) = 28.363, *p* < .001, *η*^2^ = .191]. Consistent with button RT results, post hoc tests revealed that unlike the simple ST in Experiment 1, ST Go trial EMG RTs were not significantly slower than EMG RTs from the GT, indicating participants did not slow in anticipation of stopping [*F* (1,27) = 1.32, *p =* .260, *η*^2^ = .047]. EMG RTs were quicker for the FDI compared with ADM [*F* (1,27) = 22.27, *p* < .001, *η*^2^ = .075]. As expected, failed stop trial EMG RTs had the shortest latencies in both fingers. Participants exhibited a similar amount of prEMG on successful stop trials as in Experiment 1 and contributed on average 17.7 ± 3.0 successful stop trials with prEMG per finger.

EMDs (mean ± sd) were calculated for GT go trials (FDI: 195 ± 46 ms, ADM: 186 ± 43 ms), ST go trials (FDI: 193 ± 29 ms, ADM: 177 ± 36 ms), and failed stop trials (FDI: 165 ± 27 ms, ADM: 156 ± 29 ms). Differences were observed between fingers [*F* (1,27) = 6.491, *p =* .017, *η*^2^ = .023] and across conditions [*F* (2,27) = 27.013, *p* < .001, *η*^2^ = .116]. A post-hoc 2 × 2 repeated measures ANOVA of EMDs revealed significant differences between ST go and failed stop trials [*F* (1,27) = 55.67, *p* < .001, *η*^2^ = .135, Bonferroni-corrected] and between fingers [*F* (1,27) = 8.178, *p* < 0.01, *η*^2^ = .036], but no interaction. In other words, the time between muscle activation and the button press differed between failed stop and go trials in addition to the earlier EMG onset time for failed stop than ST go trials. This suggests that there may be two distinct epochs that contribute to failed stopping: one that is prior to muscle engagement and one that occurs between muscle engagement and response completion.

#### Experiment 2: Response EMG Profiles

As above, EMG onset to EMG peak, EMG peak to EMG offset, and cancel time were calculated for all possible response conditions (Figure 3 A & C). Cancel time did not differ between fingers (*p =* .38, *d* = .17). Analysis of EMG onset to peak showed a significant main effect of finger [*F* (1,27) = 5.44, *p <* .05, *η*^2^ = .034], with longer onset to peak durations for FDI than ADM in all response conditions. There was also a main effect of response condition [*F* (3,27) = 13.25, *p* < .001, *η*^2^ = .78] with GT and ST Go responses showing longer onset to peak times than failed stop and prEMG responses. EMG peak to offset time differences were observed between fingers [*F* (1,27) = 5.513, *p* < .05, *η*^2^ = .038] and across conditions [*F* (3,27) = 5.02, *p <* .05, *η*^2^ = .025]. A significant interaction was present between finger and trial type [*F* (3,81) = 4.69, *p <* .05, *η*^2^ = .006], explained by greater peak to offset times in the GT than ST go trials for the FDI, but the reverse pattern for ADM peak to offset times for the same trial types (Figure 4 B & D). Like Experiment 1, a Bonferroni-corrected post-hoc 2 × 2 repeated measures ANOVA revealed peak to offset times were significantly shorter for failed stop than ST go trials [*F* (1,27) = 17.64, *p* < .001, *η*^2^ = .013] and that ADM peak to offset times were significantly shorter than FDI [*F* (1,27) = 6.38, *p =* .018, *η*^2^ = .063] likely due to properties of EMG recording. This is likely attributable to larger amplitude EMG recordings for FDI than ADM. Additionally, the derived slope from peak EMG to EMG offset was steeper for FDI trials compared with ADM [*F* (1,27) = 9.24, *p =* .005, *η*^2^ = .028]. Notably, failed stop trials had significantly steeper EMG peak to offset slopes compared with go trials and prEMG trials [*F* (2,27) = 56.09, *p* < .001, *η*^2^ = .252]. Consistent with Experiment 1, this result suggests the influence of a persistent active inhibitory process during failed stopping.

**Figure 3.**
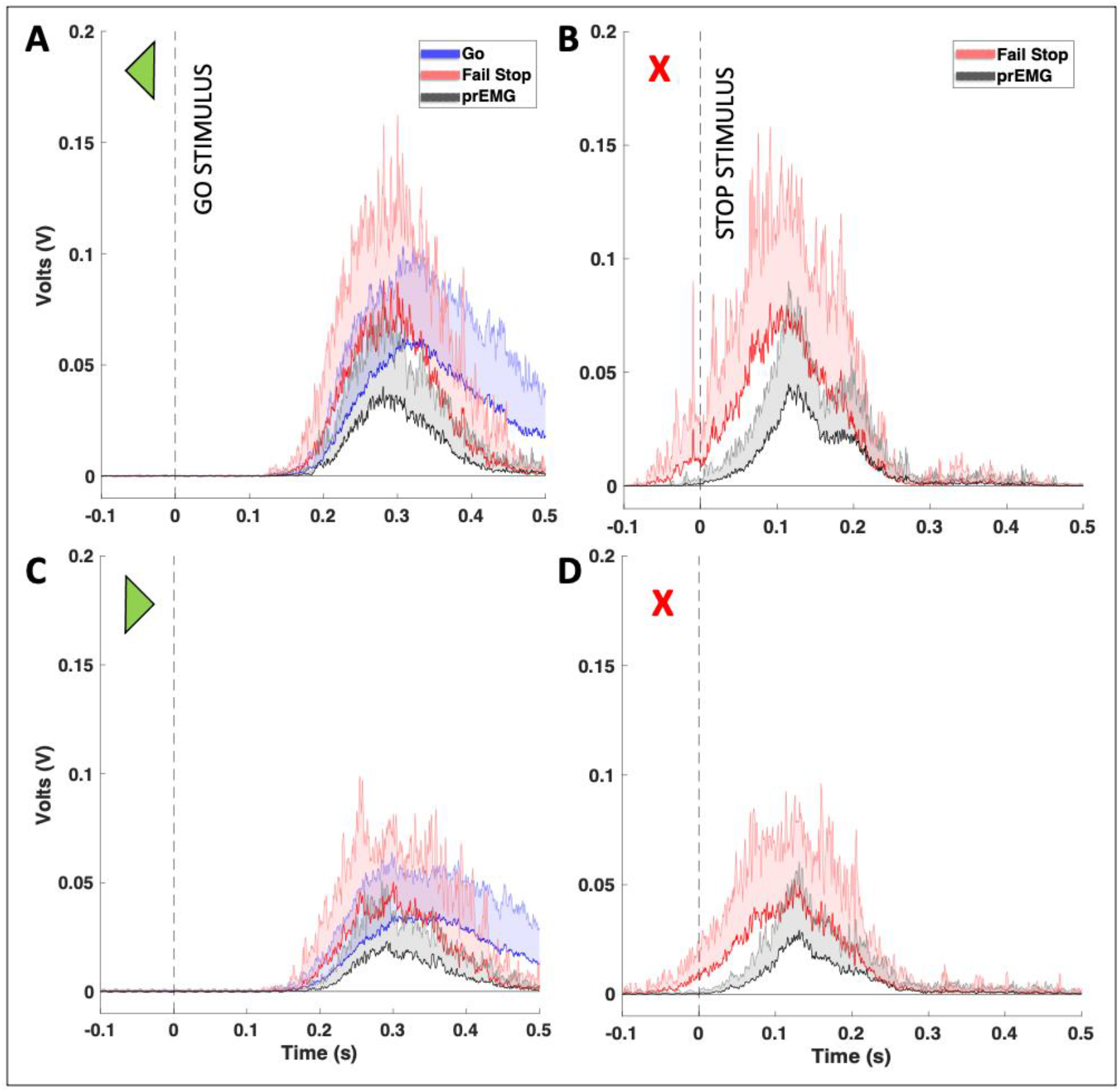
Experiment 2 (n=28) mean ± std response EMG aligned with Go stimulus (A & C) and Stop stimulus (B & D). Responses from the right FDI are in A & B, while responses from the right ADM are in C & D. Blue traces are from go trials, while black and red traces are from successful stop trials that had a partial response (prEMG) and failed stop trials, respectively. Data are smoothed via root mean square method and rectified.

**Figure 4.**
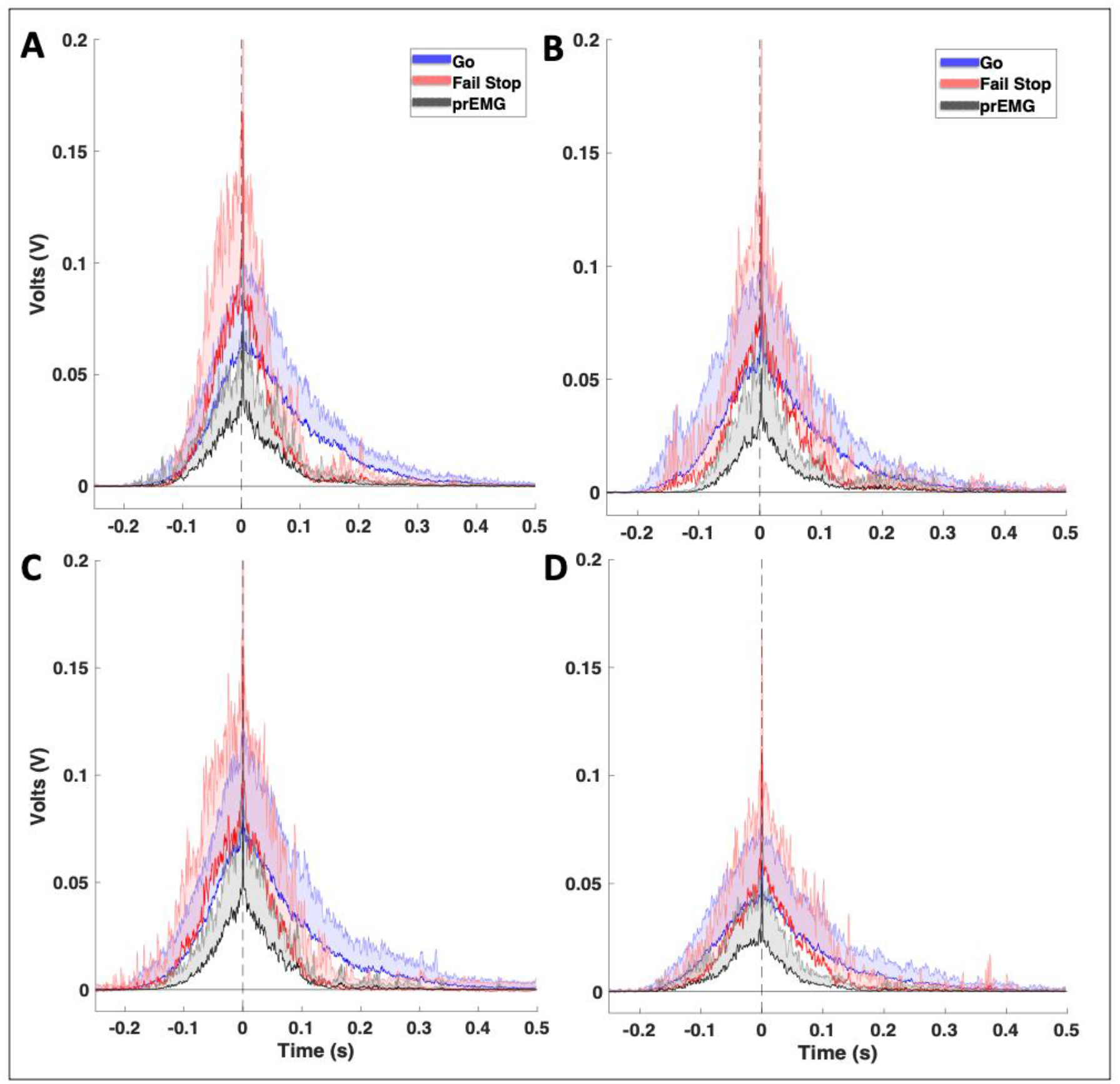
Experiment 1 (n=24) right hand (A) and left hand (C), and Experiment 2 (n=28) right FDI (B) and right ADM (D) mean ± std response EMG aligned to the EMG peak. Blue traces are from go trials, while black and red traces are from successful stop trials that had a partial response (prEMG) and failed stop trials, respectively. Data are smoothed via root mean square method and rectified.

Peak EMG (mean ± sd) values were as follows: GT go trials (FDI: 1.82 ± 0.80 V, ADM: 1.15 ± 0.63 V), ST go trials (FDI: 1.84 ± 0.79 V, ADM: 1.17 ± 0.61 V), failed stop trials (FDI: 1.81 ± 0.80 V, ADM: 1.14 ± 0.56 V) and prEMG trials (FDI: 1.02 ± 0.50 V, ADM: 0.63 ± 0.33 V). Significant differences were observed between fingers [*F* (1,27) = 25.79, *p* < .001, *η*^2^ = .157] as well as across conditions [*F* (3,27) = 90.66, *p* < .001, *η*^2^ = .145]. An interaction was observed between finger and trial type [*F* (3,81) = 4.75, *p =* .004, *η*^2^ = .006] reflecting the proportionally smaller prEMG peak amplitudes for FDI than ADM.

As for Experiment 1, we compared rectified mean EMG amplitude within the 100 ms before the stop signal between failed stop and prEMG trials (Figure 3 B & D). Paired-samples t-tests revealed that EMG activity preceding the stop signal on failed stop trials was greater than prEMG trials for both FDI and ADM (*p* < .05, d’s > .06). This replicated the findings of Experiment 1 and again indicates that EMG activity prior to the stop signal differed between successful and failed stop trials.

## DISCUSSION

In this study, we used simple and choice response versions of the stop signal task paired with EMG to investigate whether muscle activity prior to the stop signal differentiates between successful and failed stopping and whether the decline in EMG activity at the culmination of a response reflects inhibitory processes acting at the level of the muscle. We found that EMG activity in the responding finger differentiated successful and failed stopping within the 100 ms preceding the stop signal. This revisits questions partially addressed in previous studies regarding successful stop trials with partial EMG responses (de Jong et al., 1990; Jana; Raud et al., 2017). However, here we show that this difference emerges within the first 100ms prior to the stop signal in both the Simple and Choice stop tasks (Figures 2 & 3). We also describe a novel marker of inhibition in the motor system when stopping fails. The time from the EMG peak to offset was consistently shorter for failed stop than go responses. This pattern is somewhat surprising and suggests that a response inhibition mechanism continues to exert an influence on motor activity throughout response execution.

An important feature of the study was the similarity between the two experiments which allowed for relative comparisons between effectors and made it possible to assess how introducing a choice element impacts elements of task performance and recruitment of muscle activity. The design afforded opportunities for internal replication and a rich dataset of EMG measures. Each experiment provided sufficient trial numbers in a relatively large sample size to measure partial responses, and the simple and choice task results indicate consistent findings. This experimental design offered high temporal resolution and precision enabling a detailed analysis of EMG signatures between conditions. The behavioral measurements were very well matched between response hands and effectors indicating tasks were performed similarly and outcomes were comparable. However, slowing was observed only in the simple version of the stop task (Experiment 1). The lack of slowing in Experiment 2 compared to Experiment 1 may be a consequence of the involvement of a choice, which inherently lengthens RTs. With our two separate samples, we cannot sufficiently address this question as this pattern may also be the consequence of inter-individual differences. Nevertheless, the presence or absence of slowing did not appear to affect our EMG measures of interest related to stopping.

Previous work in this area (Jana et al., 2020; Raud et al., 2017) investigated aspects of EMG markers with particular focus on trials with partial responses. However, they did not examine the extent and timing of EMG activation as a determinant of behavioral performance. Inconsistent with earlier work investigating the “point of no return” (De Jong et al., 1990), we observed that EMG increases prior to the stop signal led to stopping failure, regardless of whether a response choice was involved. This may be in line with more recent evidence indicating stopping is not faster than going (Du et al., 2022) as is implicit to the traditional horse race model. Our data provides physiological evidence that successful cancellation of a response may not be possible if EMG activity is present prior to the stop signal.

We also found new evidence indicating the inhibitory process continues even when stopping fails. Our observation that EMG ramps down more quickly when stopping is unsuccessful provides a novel physiological marker of inhibition acting on the responding muscle. This inhibition may influence ensuing behavior on subsequent trials. An interesting question concerns whether or not this marker can be detected using other versions of the stop signal task, such as the anticipatory response inhibition task, to potentially discriminate between selective and non-selective inhibitory mechanisms (MacDonald et al., 2014; Wadsley, Cirillo, & Byblow, 2019).This marker may be sensitive to abnormalities in inhibition in clinical populations such as Parkinson’s Disease (Gaugel et al., 2004; Swann et al., 2011; Obeso et al., 2008; MacDonald & Byblow, 2015). Furthermore, stopping may depend on two inhibitory mechanisms that act in sequence to pause-then-cancel a response (Frank, 2006; Schmidt & Berke, 2017; Diesburg & Wessel, 2021), and the lingering inhibition we observed during failed stopping may be a distinct marker of the second part of this two-step stopping process. These signatures may also have value for differentiating stopping related inhibitory processes from other markers of motor inhibition, for example those associated with unexpected and infrequent events (Iacullo, Diesburg, & Wessel, 2020).

### Limitations

A limitation of this study was the notably high success rate at stopping in both experiments, above the typical 50% target. While a high rate of stopping can invalidate SSRT estimates, here the higher rate of successful stopping afforded a greater number of partial EMG responses. This was helpful for attaining sufficient statistical power to compare EMG activity across response conditions. Additionally, the Simple stop task design in Experiment 1 deviates from the recommended guidelines for administration of the stop task (Verbruggen et al., 2019) because the go task does not include a response choice. We specifically selected this design to assess response initiation in the absence of a choice decision, and the convergent results between the two experiments provide validation that effects are not the result of interactions with response choice decision-related processes.

Another possible limitation is that the tasks only included visual stimuli, which could leave open possible alternative outcomes for other modalities and warrants further investigation. Finally, the design of each experiment included a tonic contraction that was held with the non-responding hand for the duration of data collection. This decision was motivated by our initial intention to examine the tonic signal for EMG signatures of widespread inhibition when participants successfully canceled movements, in line with TMS experiments of a similar design (Badry et al., 2009; Cai et al., 2012; Majid et al., 2012; Wessel et al., 2013b). Our initial approach to the analysis did not reveal stopping-related modulation within the tonic EMG signal, and we chose not to present these data here as follow-up investigations are underway. The additional task demands associated with sustaining a muscle contraction at a fixed level may have influenced stop task performance. However, this does not seem likely as RTs and SSRT values are comparable to studies with similar task designs.

## Conclusions

Previous work has not fully addressed whether a point of no return exists and whether inhibitory processes associated with stopping persist even during action execution. We addressed these questions in two experiments with a close examination of EMG activity. EMG activity preceding the stop signal differentiated successful and failed stop trials. This strongly supports the existence of a point of no return prior to the stop signal. Nevertheless, partial EMG responses on successful stop trials indicate that the stop process runs to completion even after the responding muscle is engaged. Additionally, the rate of decline in EMG activity on failed stop trials suggested a lingering effect of inhibition in the motor system. These two non-invasive electrophysiological markers associated with stop task performance may be useful for isolating specific mechanisms engaged in reactive stopping and may help to improve our understanding of inhibitory control deficits in clinical populations.

## Acknowledgements

We would like to thank Isabella Utrecht and Chris Horton for assistance with data collection and analysis, Nick Jackson for designing and constructing the response devices, and Nicole Swann and Kelsey Schultz for feedback on the manuscript.

## REFERENCES

Badry, R., Mima, T., Aso, T., Nakatsuka, M., Abe, M., Fathi, D., Foly, N., Nagiub, H., Nagamine, T., & Fukuyama, H. (2009). Suppression of human cortico-motoneuronal excitability during the stop-signal task. Clinical Neurophysiology, 120, 1717–1723. https://doi.org/10.1016/j.clinph.2009.06.027

Bissett, P. G., Jones, H. M., Poldrack, R. A., & Logan, G. D. (2021). Severe violations of independence in response inhibition tasks. Science advances, 7(12). https://doi.org/10.1126/sciadv.abf4355

Cai, W., Oldenkamp, C. L., & Aron, A. R. (2012). STopping speech suppresses the task-irrelevant hand. Brain and language, 120, 412–415. https://doi.org/10.1016/j.bandl.2011.11.006

De Jong, R., Coles, M. G. H., Logan, G. D., & Gratton, G. (1990). In search of the point of no return: The control of response process. Journal of Experimental Psychology: Human Perception and Performance, 16, 164–181. https://doi.org/10.1037/0096-1523.16.1.164

Diesburg, D. A., & Wessel, J. R. (2021). The Pause-then-Cancel model of human action-stopping: theoretical considerations and empirical evidence. Neuroscience & Biobehavioral Reviews, 129, 17–34. https://doi.org/10.1016/j.neubiorev.2021.07.019

Du, Y., Forrence, A. D., Metcalf, D. M., & Haith, A. M. (2022). Action inhibition revisited: Stopping is not faster than going. bioRxiv. https://doi.org/10.1101/2022.06.29.497798

Frank, M. J. (2006). Hold your horses: a dynamic computational role for the subthalamic nucleus in decision making. Neural networks, 19(8), 1120–1136. https://doi.org/10.1016/j.neunet.2006.03.006

Gauggel, S., Rieger, M., & Feghoff, T. (2004). Inhibition of ongoing responses in patients with Parkinson’s disease. Journal of Neurology, Neurosurgery & Psychiatry, 75(4), 539–544. https://doi.org/10.1136/jnnp.2003.016469

Goonetilleke, S. C., Doherty, T. J., & Corneil, B. D. (2010). A within trial measure of the stop signal reaction time in a head-unrestrained oculomotor countermanding task. Journal of neurophysiology, 104(6), 3677–3690. https://doi.org/10.1152/jn.00495.2010

Iacullo, C., Diesburg, D. A., & Wessel, J. R. (2020). Non-selective inhibition of the motor system following unexpected and expected infrequent events. Experimental brain research, 238(12), 2701–2710. https://doi.org/10.1007/s00221-020-05919-3

Jackson, N., & Greenhouse, I. (2019). VETA: an open-source matlab-based toolbox for the collection and analysis of electromyography combined with transcranial magnetic stimulation. Frontiers in Neuroscience, 13, 975. https://doi.org/10.3389/fnins.2019.00975

The jamovi project (2021). jamovi. (Version 2.2) [Computer Software]. Retrieved from https://www.jamovi.org.

Jana, S., Hannah, R., Muralidharan, V., & Aron, A. R. (2020). Temporal cascade of frontal, motor and muscle processes underlying human action-stopping. Elife, 9, e50371. https://doi.org/10.7554/eLife.50371

JASP Team. (2020). JASP (Version 0.14.1)[Computer software]. 2020. URL https://jasp-stats.org.

Logan, G. D., & Cowan, W. B.(1984). On the ability to inhibit thought and action: A theory of an act of control. Psychological Review, 91, 295–327. https://doi.org/10.1037/0033-295X.91.3.295

Macdonald, H. J., Coxon, J. P., Stinear, C. M., & Byblow, W. D. (2014). The fall and rise of corticomotor excitability with cancellation and reinitiation of prepared action. Journal of neurophysiology, 112(11), 2707–2717. https://doi.org/10.1152/jn.00366.2014

MacDonald, H. J., & Byblow, W. D. (2015). Does response inhibition have pre-and postdiagnostic utility in Parkinson’s disease?. Journal of motor behavior, 47(1), 29–45. https://doi.org/10.1007/s00221-020-05919-3

Majid, D. S. A., Cai, W., George, J. S., Verbruggen, F., & Aron, A. R. (2012). Transcranial magnetic stimulation reveals dissociable mechanisms for global versus selective corticomotor suppression underlying the stopping of action. Cerebral Cortex, 22, 363–371. https://doi.org/10.1093/cercor/bhr112

Matzke, D., Curley, S., Gong, C. Q., & Heathcote, A. (2019). Inhibiting responses to difficult choices. Journal of Experimental Psychology: General, 148(1), 124. https://doi.org/10.1037/xge0000525

Matzke, D., Love, J., & Heathcote, A. (2017). A Bayesian approach for estimating the probability of trigger failures in the stop-signal paradigm. Behavior research methods, 49(1), 267–281. https://doi.org/10.3758/s13428-015-0695-8

McGarry, T., Inglis, J. T., & Franks, I. M. (2000). Against a final ballistic process in the control of voluntary action: evidence using the Hoffmann reflex. Motor control, 4(4), 469–485. https://doi.org/10.1123/mcj.4.4.469

Obeso, J. A., Rodríguez-Oroz, M. C., Benitez-Temino, B., Blesa, F. J., Guridi, J., Marin, C., & Rodriguez, M. (2008). Functional organization of the basal ganglia: therapeutic implications for Parkinson’s disease. Movement disorders: official journal of the Movement Disorder Society, 23(S3), S548–S559. https://doi.org/10.1002/mds.22062

Raud, L., Thunberg, C., & Huster, R. J. (2022). Partial response electromyography as a marker of action stopping. Elife, 11, e70332. https://doi.org/10.7554/eLife.70332

Raud, L., Huster, R.J., Ivry, R.B., Labruna, L., Messel, M.S., & Greenhouse, I. (2020). A single mechanism for global and selective response inhibition under the influence of motor preparation. Journal of Neuroscience, 40(41). https://doi.org/10.1523/JNEUROSCI.0607-20.2020

Raud, L., & Huster, R. J. (2017). The temporal dynamics of response inhibition and their modulation by cognitive control. Brain topography, 30(4), 486–501. https://doi.org/10.1007/s10548-017-0566-y

Schmidt, R., & Berke, J. D. (2017). A Pause-then-Cancel model of stopping: evidence from basal ganglia neurophysiology. Philosophical Transactions of the Royal Society B: Biological Sciences, 372(1718), 20160202. https://doi.org/10.1098/rstb.2016.0202

Skippen, P., Matzke, D., Heathcote, A., Fulham, W. R., Michie, P., & Karayanidis, F. (2019). Reliability of triggering inhibitory process is a better predictor of impulsivity than SSRT. Acta psychologica, 192, 104–117. https://doi.org/10.1016/j.actpsy.2018.10.016

Swann, N., Poizner, H., Houser, M., Gould, S., Greenhouse, I., Cai, W., … & Aron, A. R. (2011). Deep brain stimulation of the subthalamic nucleus alters the cortical profile of response inhibition in the beta frequency band: a scalp EEG study in Parkinson’s disease. Journal of Neuroscience, 31(15), 5721–5729. https://doi.org/10.1523/JNEUROSCI.6135-10.2011

Verbruggen, F., & Logan, G. D. (2009). Proactive adjustments of response strategies in the stop-signal paradigm. Journal of Experimental Psychology: Human Perception and Performance, 35(3), 835. https://psycnet.apa.org/doi/10.1037/a0012726

Verbruggen, F., Aron, A. R., Band, G. P., Beste, C., Bissett, P. G., Brockett, A. T., … & Boehler, C. N. (2019). A consensus guide to capturing the ability to inhibit actions and impulsive behaviors in the stop-signal task. elife, 8, e46323. https://doi.org/10.7554/eLife.46323.027

Wadsley, C. G., Cirillo, J., & Byblow, W. D. (2019). Between-hand coupling during response inhibition. Journal of neurophysiology, 122(4), 1357–1366. https://doi.org/10.1152/jn.00310.2019

Wessel, J. R., & Aron, A. R. (2017). On the globality of motor suppression: unexpected events and their influence on behavior and cognition. Neuron, 93(2), 259–280. https://doi.org/10.1016/j.neuron.2016.12.013

Wessel, J. R., Ghahremani, A., Udupa, K., Saha, U., Kalia, S. K., Hodaie, M., … & Chen, R. (2016). Stop-related subthalamic beta activity indexes global motor suppression in Parkinson’s disease. Movement Disorders, 31(12), 1846–1853. https://doi.org/10.1002/mds.26732

Wessel, J. R., & Aron, A. R. (2013). Unexpected events induce motor slowing via a brain mechanism for action-stopping with global suppressive effects. Journal of Neuroscience, 33(47), 18481–18491. https://doi.org/10.1523/JNEUROSCI.3456-13.2013

Wessel, J. R., Reynoso, H. S., & Aron, A. R. (2013b). Saccade suppression exerts global effects on the motor system. Journal of Neurophysiology, 110, 883–890. https://doi.org/10.1152/jn.00229.2013

